# A spatiotemporal complexity architecture of human brain activity

**DOI:** 10.1101/2021.06.04.446948

**Authors:** Stephan Krohn, Nina von Schwanenflug, Leonhard Waschke, Amy Romanello, Martin Gell, Douglas D. Garrett, Carsten Finke

**Affiliations:** Department of Neurology, Charité-Universitätsmedizin Berlin; Berlin, Germany; Berlin School of Mind and Brain, Humboldt-Universität zu Berlin; Berlin, Germany; Center for Lifespan Psychology, Max Planck Institute for Human Development; Berlin, Germany; Max Planck UCL Centre for Computational Psychiatry and Ageing Research; Berlin, Germany; Institute of Neuroscience and Medicine (INM-7: Brain & Behavior), Research Centre Jülich, Jülich, Germany; Department of Psychiatry, Psychotherapy and Psychosomatic Medicine, RWTH Aachen University, Aachen, Germany

## Abstract

The human brain operates in large-scale functional networks. These networks are thought to arise from neural variability, yet the principles behind this link remain unknown. Here we report a mechanism by which the brain’s network architecture is tightly linked to critical episodes of neural regularity, visible as spontaneous ‘complexity drops’ in functional MRI signals. These episodes support the formation of functional connections between brain regions, subserve the propagation of neural activity, and reflect inter-individual differences in age and behavior. Furthermore, complexity drops define neural states that dynamically shape the coupling strength, topological structure, and hierarchy of brain networks and comprehensively explain known structure-function relationships within the brain. These findings delineate a unifying complexity architecture of neural activity – a human ‘complexome’ that underpins the brain’s functional network organization.

---

The human brain operates in large-scale functional networks that underpin cognition and behavior^1–7^. Collectively subsumed as the functional human *connectome*^8^, these networks are an expression of temporally correlated activity across spatially distributed brain regions. This correlation structure is commonly referred to as functional connectivity (FC)^9–11^, and the estimation of FC from functional magnetic resonance imaging (fMRI) has revealed important insights into the brain’s functional architecture. First, FC is not uniformly distributed across the brain, but rather organized in functional subsystems known as resting-state networks (RSN)^1,2,4,6,12^. Second, FC is not static over time, but shows dynamic fluctuations that result in distinct temporal network states^13–17^. Third, the structure of the network is not random, but follows an efficient topological configuration subserving network economy^18,19^. Finally, brain networks are not functionally equivalent, but are organized along a principal hierarchy spanning from lower-order unimodal to higher-order transmodal processing systems^20–22^.

Despite these advances, it remains a fundamental challenge in neuroscience to understand what determines the structure, temporal dynamics, and hierarchy within the human connectome. To achieve such an understanding, an explanatory framework is required that links these global network properties to the local activity of individual regions. Such a link is crucial because functional connectivity – and all network properties derived from it – are defined on the relationship *between* regional signals. Given this unidirectionality of network construction (regional activity defines inter-regional connections, but not vice versa), the emergence of functional networks must ultimately be rooted in these signals themselves. To this effect, the intrinsic variability of neural activity fluctuations is thought to facilitate network formation^4,23– 26^, but how variability enables connectivity has not been shown. In this regard, recent work on temporal brain dynamics suggests that FC may arise from only a few decisive episodes in the blood oxygen level-dependent (BOLD) brain signals^16,27–29^, yet the nature of such critical moments remains unclear.

## Results

### Brain activity is characterized by critical moments of neural regularity

To close this gap, we here employ an information-theoretic complexity analysis that (i) relates the local activity of individual brain regions to the global properties of the network, (ii) represents the intrinsic variability of neural fluctuations in a standardized space, and (iii) allows for a time-resolved account of neural dynamics to capture critical moments in the signals. This approach rests on symbolic encoding of BOLD activity and leverages both amplitude information and the diversity of abstract patterns in the signal to quantify the degree of irregularity (i.e., signal complexity) over a given moment in time^30^ (Extended Data Fig. 1-2). We thus obtain a complexity timeseries for each brain region, which captures local dynamics through a meta-representation of neural variability and allows for a link between the nodes in the network (i.e., individual regions) and the edges in the network (i.e., region-to-region functional connections).

Application of this approach to resting-state fMRI from the Human Connectome Project (HCP)^8^ revealed a highly canonical complexity structure of human brain activity: Over ∼80% of the acquisition time, the brain exhibits widespread high-complexity signals (HCS), approaching the upper limit of expected values given the underlying signal characteristics (Extended Data Fig. 1-3). Intermittently, however, these HCS are interrupted by spontaneous ‘complexity drops’ that correspond to episodes of increased regularity in the BOLD signal (Fig. 1a, inset). These complexity drops appear in clusters of brain regions and over well-delineated moments in time, and the number of brain regions engaging in them closely reflects the current overall complexity of neural activity. Notably, this complexity structure was universally present in all analyzed recordings (684 scans from n=343 participants; also see online repository for individual scans, and validation in holdout data) and was highly robust against different windowing parameters, parcellation granularity, and timeseries extraction methods (Extended Data Fig. 4-5).

**Fig. 1.**
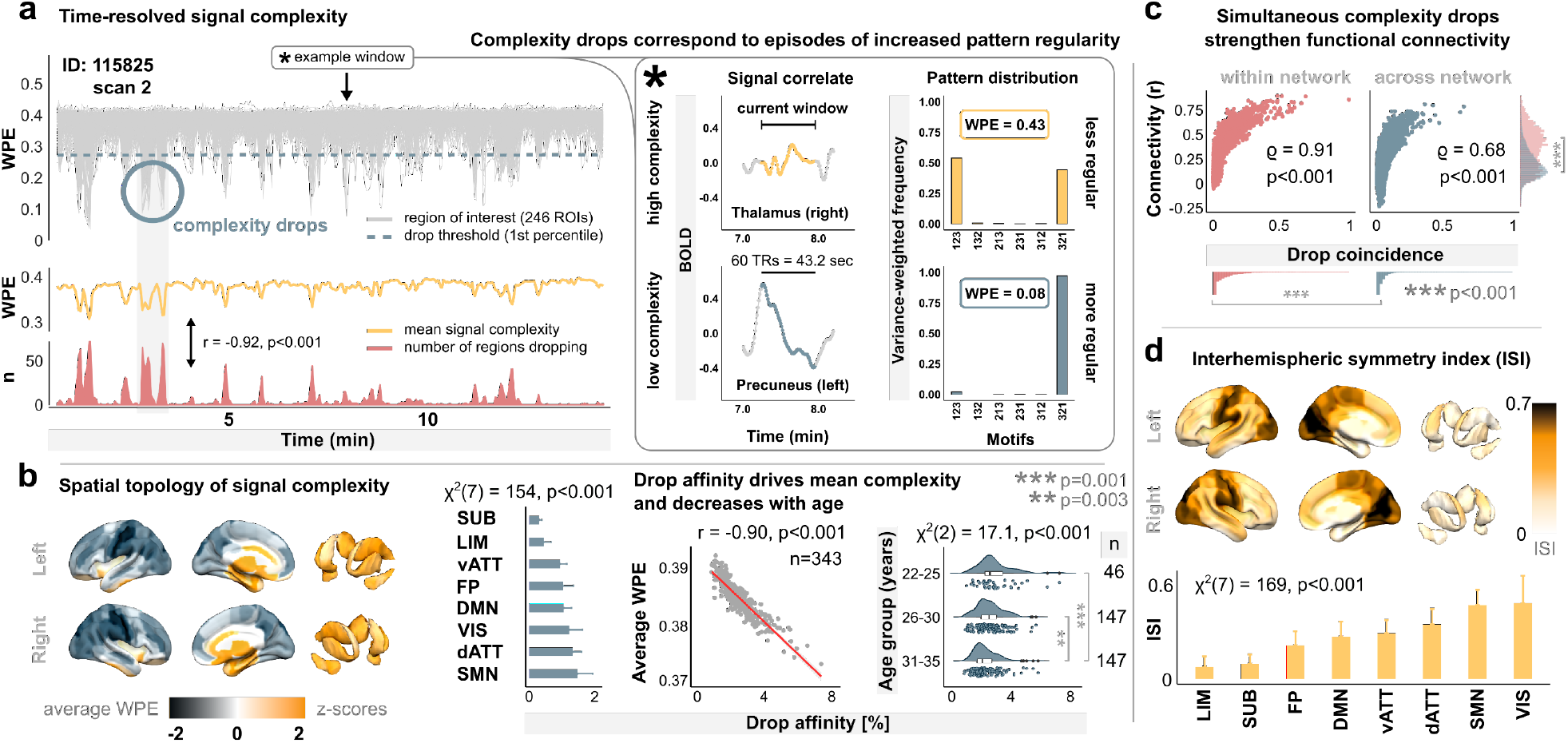
Human brain activity is characterized by critical moments of neural regularity that are visible as transient ‘complexity drops’. **a**, Time-resolved weighted permutation entropy (WPE) on the neural signals from each region of interest (ROI) in an exemplary resting-state recording (Brainnetome parcellation; windows length 60 TRs, 95% overlap). The inset displays the BOLD correlates of two representative high- and low-complexity signals. **b**, Spatial distribution of average signal complexity. Drop affinity over RSN, link to grand-average signal complexity, and age-related reduction in drop affinity. **c**, FC as a function of drop coincidence, within and across canonical RSN. **d**, Spatial topology, and network distribution of interhemispheric complexity symmetry. Network abbreviations: dATT: dorsal attention; DMN: default mode; FP: frontoparietal; LIM: limbic; SMN: somatomotor; SUB: subcortical; vATT: ventral attention; VIS: visual.

Spatial analysis revealed persistent HCS in medial temporal, anterior cingulate and subcortical regions – most markedly thalamic areas (Fig. 1b). In contrast, complexity drops predominantly occurred in the cortex, with a focus on pericentral regions. On the level of individual participants, the affinity for complexity drops was strongly linked to grand-average brain signal complexity (r=-0.9, p=1.6e-125) and showed significant differences across RSN (χ^2^(7)=154, p=6.1e-30) as well as gradual decreases with age (χ^2^(2)=17.1, p=1.9e-4), even over the narrow age range of early adulthood (Fig. 1b).

This time-resolved representation of BOLD dynamics subsequently allowed us to relate the complexity timeseries of individual regions to the functional coupling between those regions. Analyzing the co-occurrence of complexity drops across the brain revealed that FC strength between any two regions is strongly associated with the degree to which they engage in complexity drops together (Fig. 1c). While this ‘drop coincidence’ explained a large part of the variance in FC overall (F(1,30133)=4.3e4, p≈0, R^2^_adj_=0.59), it was also significantly higher for connections within, rather than across canonical RSN (W=3.6e7, p=9.3e-161), paralleled by higher within-than across-network connectivity.

A further link to such network differentiation is demonstrated by the interhemispheric symmetry of regional complexity timeseries (Fig. 1d), quantified as the rank-weighted correlation of each region with its contralateral equivalent. This interhemispheric symmetry was systematically constrained by canonical RSN (χ^2^(7)=169, p=4.6e-33) and intrinsically mirrored the recently uncovered hierarchy between unimodal and transmodal areas^20,21^, suggesting that the cross-hemispheric coupling of neural activity patterns is strongest in primary networks and becomes gradually more diverse in higher-order systems.

### Moments of regularity spread throughout the brain

Recent work has suggested a propagation of BOLD activity patterns throughout the brain as a means of communication within the network^31–33^. As the symbolic encoding approach used here inherently captures such neural patterns^30^, we asked if complexity drops may subserve the propagation of neural activity across the brain. Supporting this idea, complexity drops consistently occurred in dynamic cascades that start with a few initializing regions, gradually spread across the brain, and finally fade back to a few regions again, such that brain-wide drop engagement exhibits an inverted U-shape over time (Fig. 2a; Supplementary Movie 1). Such cascades were present in all 343 participants and typically lasted about 10 seconds (mean duration 9.4 ± 3.5s, range 6.5 – 41s). To analyze the spread of complexity drops across the brain, we developed a graph theoretical framework, where each individual propagation is described by a directed graph (Fig. 2b). These propagation graphs contain node layers that represent consecutive BOLD windows from initialization of the cascade (‘source layer’) to maximum drop engagement (‘peak layer’), and where the direction of edges expresses progression in time, allowing for the estimation of region-to-region transition probabilities within the propagation.

**Fig. 2.**
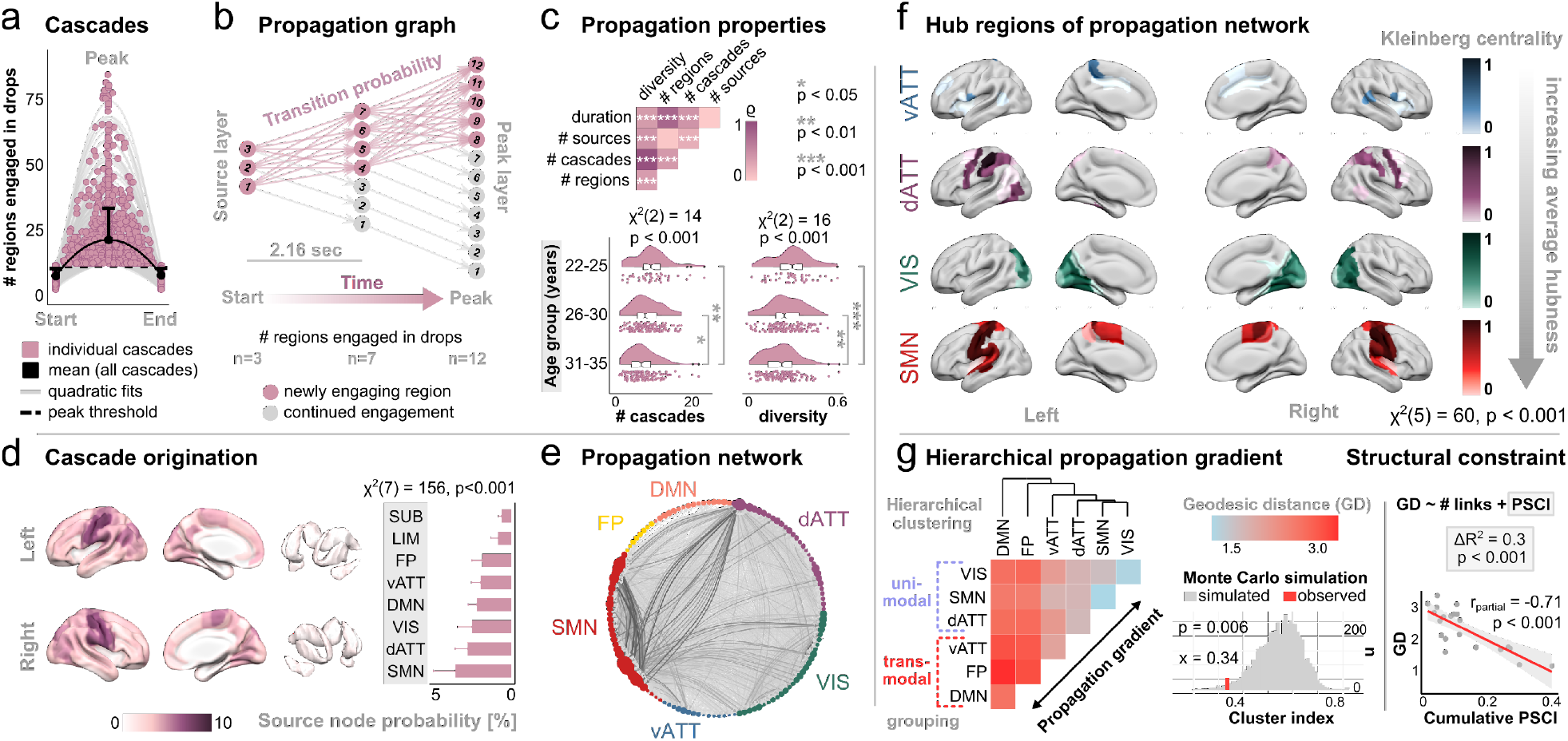
Complexity drops spread throughout the brain along a principal functional hierarchy. **a**, Drop cascades. Points represent a random sample of 500 out of 5279 total cascades in the study population. Engagement is thresholded to at least 10 regions dropping simultaneously at peak. **b**, Formalization of individual spreads as directed propagation graphs. Window-to-window temporal resolution corresponds to 2.16 seconds (3 TRs). **c**, Propagation properties and age effects (pairwise tests FDR-corrected). **d**, Spatial topology of cascade origination. **e**, Propagation network, thresholded to the 95th percentile of strongest connections (edge thickness: transition probability; node diameter: source probability). **f**, Hub topology of the propagation network, given by Kleinberg’s eigenvector centrality. FP and DMN are omitted because of near-zero entries only. **g**, Average node-to-node geodesic distances in the propagation network. Monte Carlo simulation testing the significance of unimodal vs. transmodal network clusters. Association between geodesic distances and the probabilistic streamline connectivity index (PSCI).

Drop cascades were characterized by a strong positive relationship between cascade duration and the number of regions engaging in them (ρ=0.79, p_adj_=2.9e-73), and the number of cascades an individual presented was strongly related to their source diversity, defined as the proportion of unique brain regions ever initializing a cascade (ρ=0.96, p_adj_=2e-182; Fig. 2c). Notably, we observed significant age-related reductions in the number of cascades (χ^2^(2)=13.9, p=9.5e-4) and in source diversity (χ^2^(2)=16.3, p=2.8e-4), but not in the number of regions dropping (χ^2^(2)=4.5, p=0.11) or cascade duration (χ^2^(2)=2.6, p=0.27), suggesting that individual propagations occur in a semi-canonical fashion, although the predisposition to engage in them decreases with age (Fig. 2c).

Spatial analysis of propagation graphs revealed that drop cascades originate predominantly in the cortex, with a focus on lower-order RSN (Fig. 2d). Furthermore, graph construction from the transition probabilities of individual spreads uncovered a highly structured propagation network (Fig. 2e). Centrality analysis on this network revealed that the propagation of complexity drops follows an intrinsic hierarchy (Fig. 2f), where the most influential nodes correspond to primary RSN, and nodes of higher-order systems become gradually less important to the propagation.

Further supporting such a propagation hierarchy from lower-order to higher-order systems, geodesic distances in the propagation network showed that complexity drops spread along a unimodal-to-transmodal functional gradient (Fig. 2g; Monte Carlo simulation of cluster groups: x_cluster_=0.34, p=0.006). This propagation structure thus again strongly converged with the recently uncovered hierarchy within the connectome^20–22^, characterized by a principal gradient between lower-order unimodal and higher-order transmodal processing systems.

Furthermore, modelling geodesics as a function of probabilistic streamline connectivity and number of structural links (R^2^_adj_=0.66, F(2,18)=20.9, p=2e-5) significantly improved explanatory power (ΔR^2^=0.3, η^2^_partial_=0.5, F(1,18)=17.8, p=5.1e-4) compared to when only the number of links were considered (R^2^_adj_=0.37, F(1,19)=12.7, p=0.002), suggesting that the propagation of complexity drops is additionally constrained by structural connectivity.

### Neural activity is organized in network-modulating complexity states

Next, we related the complexity dynamics of individual regions to the behavior of the whole-brain network, testing the idea that functional states of the network^13–15^ should be rooted in underlying states of neural activity. Unsupervised clustering supported this hypothesis, showing that neural activity is organized in distinct temporal complexity states (Fig. 3a). Consistent with the inspection of individual timeseries (Fig. 1a; online repository), participants spent most time in a default neural state of high complexity, while the more infrequently visited states entailed gradually decreasing levels of complexity, yielding a strong discriminatory effect of complexity across states (η^2^=0.90, [0.89-0.91]). Notably, while complexity states are derived directly from the activity of individual brain regions, they resulted in pronounced concordant effects on the coupling strength and topological structure of the connectome, which are derived from region-to-region signal correlations. In particular, the default state of neural activity is characterized by low coupling strength of the network, while the lower-complexity states entail gradual increases in connectivity (Fig. 3a).

**Fig. 3.**
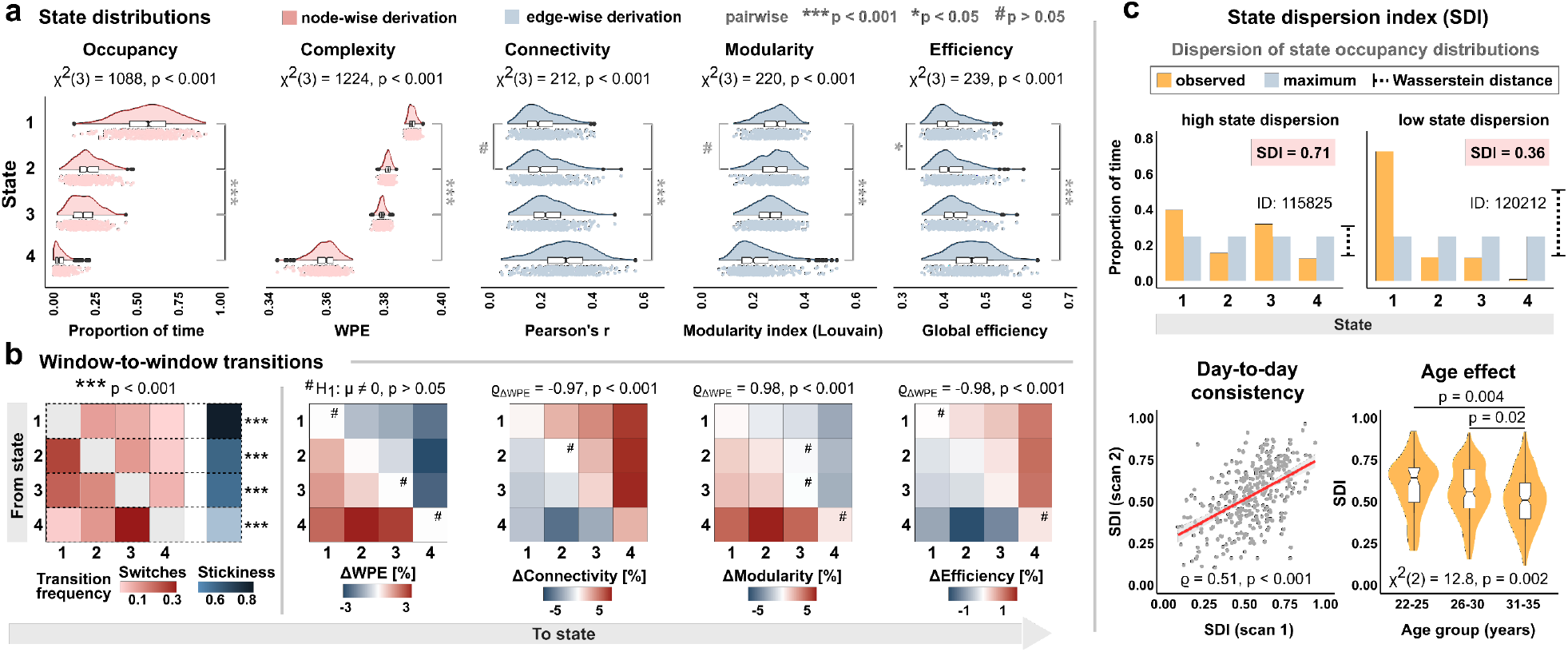
Neural activity is organized in temporal complexity states that modulate network strength and configuration. **a**, State-wise distributions of occupancy, signal complexity, coupling strength, network modularity, and global efficiency (n=343 participants). Node-wise derivation refers to estimation from individual regional signals, edge-wise derivation to estimation from the network (i.e., region-to-region correlations). **b**, Window-to-window state transition frequencies. State-dependent changes tested against deviation from zero (t-test, p_FDR_<0.05 for all except those marked by #). Association of network measures with ΔWPE corresponds to the correlation of the respective matrices. **c**, Illustration, consistency across two consecutive days, and age-related reduction of the state dispersion index (SDI). All pairwise tests FDR-corrected.

This impact of complexity states furthermore extended to the topological configuration of the connectome^5,18,19^, with the default neural state yielding high modularity and low efficiency in the network. As for FC, visiting lower-complexity states resulted in gradual decreases in modularity and concordant increases in network efficiency. Notably, this network-modulating effect of complexity states even held at the temporal resolution of individual window-to-window transitions (Fig. 3b). While all complexity states were similarly stable, switches between low-complexity and high-complexity brain states typically occurred through the intermediate states, and these transitions were accompanied by fine-grained, momentary changes in the connectivity, modularity, and efficiency of the network.

Given these group-level effects of complexity states on the network, we furthermore tested how much the space of possible complexity states was explored by individual participants (Fig. 3c). To this end, we derived a measure of state exploration based on the Wasserstein Distance between an individual’s empirical state visits and a theoretical uniform distribution. This state dispersion index (SDI) was remarkably consistent across two consecutive days of scanning (ρ=0.51, p≈0) and significantly decreased with age (χ^2^(2)=12.8, p=0.002), suggesting that older participants present increasingly rigid neural dynamics, consistent with the age-related reductions in drop affinity (Fig. 1b) and propagation diversity (Fig. 2c). Notably, all state-related findings were highly robust against the number of expected complexity states as defined by the clustering parameters (Extended Data Fig. 6-7).

### Complexity states link structural and functional network hierarchies

Next, we analyzed how these neural complexity states are spatially embedded in the brain. Given that both interhemispheric signal coupling (Fig. 1d) and the propagation of complexity drops (Fig. 2e-g) intrinsically followed the principal unimodal-to-transmodal hierarchy, we explicitly estimated the corresponding gradient loadings from the FC data and related them to the topology of complexity states. Furthermore, these gradient loadings have been shown to be spatially correlated with cortical myeloarchitecture as a proxy of anatomical hierarchy^21,34,35^, yielding an important structure-function relationship within in the connectome^36^ that also partly extends to non-primate mammalian brains^37,38^. Thus, we estimated cortical myelination as the T1-weighted/T2-weighted image ratio and related this myelin distribution to the functional gradient loadings and complexity states.

The topology of complexity states corroborated the distribution of average signal complexity (Fig. 1b), with subcortical areas persistently showing high-complexity activity, largely independent of the current global complexity state (Fig. 4a). In contrast, cortical topologies varied distinctly with the number of regions engaged in complexity drops (χ^2^(3)=1172, p=9.4e-254), inducing significant spatial heterogeneity across states (MLRT=655, p≈0). To estimate which brain regions may drive these differences, we derived a measure of across-state distance (ASD) as the cumulative centroid-to-centroid Euclidean distance for every region across 4-dimensional state space (Fig. 4b). Again, the ASD closely followed a unimodal-to-transmodal gradient, where regions that were most variable across complexity states represented the unimodal end of the hierarchy. Testing this spatial convergence with spin permutation correlation uncovered a pronounced positive association of ASD topology with cortical myeloarchitecture (ρ_empirical_=0.67, p_spin_<0.001) as well as a strong negative relationship with the connectivity gradient (ρ_empirical_=-0.76, p_spin_<0.001; Fig. 4c). While we were able to replicate the previously reported link between myelination and the FC gradient^34^ (ρ_empirical_=-0.52, p_spin_<0.001), partial correlations of all three variables revealed that the association between myelin and the gradient loadings disappears when controlling for ASD, whereas the ASD-myelin and ASD-gradient relationships persist when controlling for the respective other variable (Fig. 4d). Further corroborating this finding, a hierarchical regression approach showed that ASD topology alone explained a large part of the variance in gradient loadings (R^2^_adj_=0.57, p=4.9e-40) and that accounting for ASD in the augmented model completely resolves the myelin-gradient effect (Fig. 4d).

**Fig. 4.**
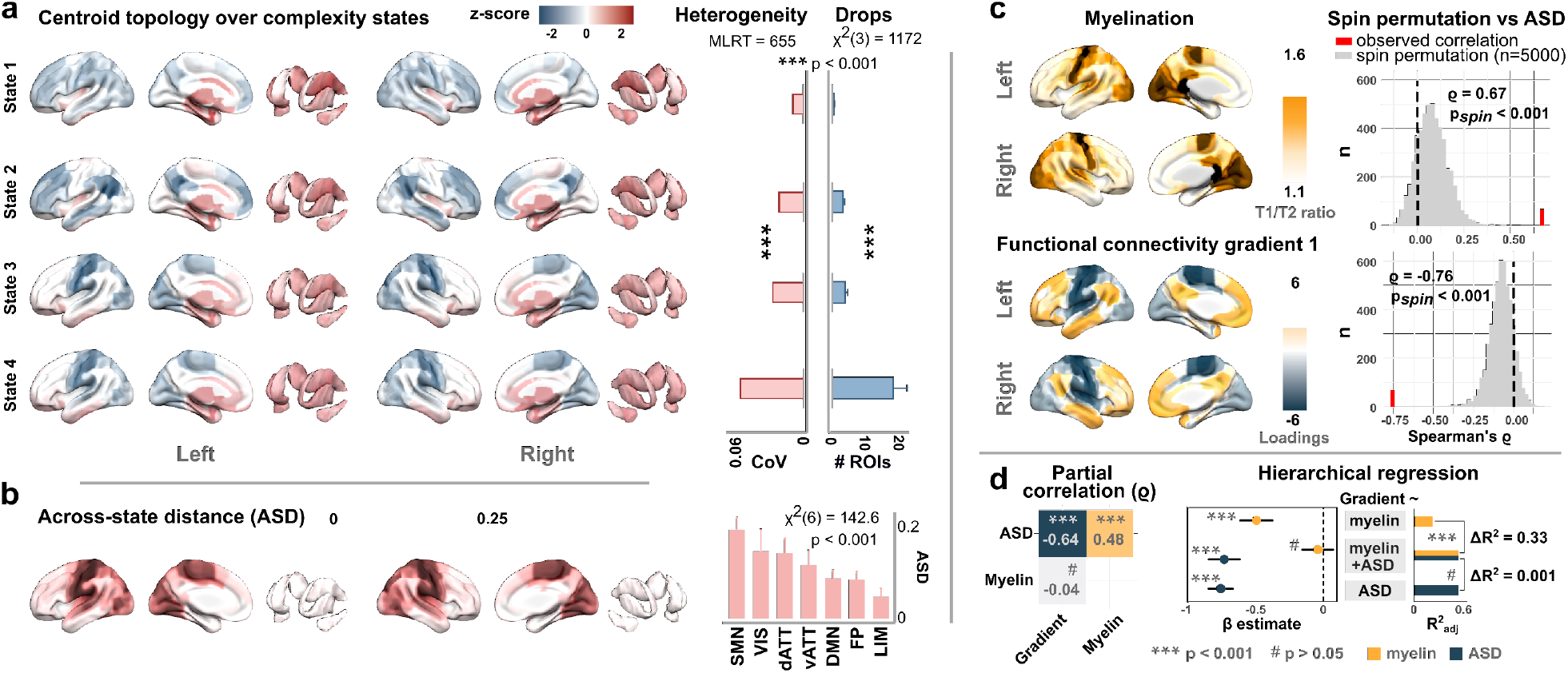
The spatial embedding of complexity states comprehensively explains structure-function relationships within the brain. **a**, Spatial topology of centroid locations (within-state z-scores). Coefficient of variation (CoV) over regions and average state-wise complexity drops (error bars: standard deviation). MLRT, modified likelihood ratio test. **b**, Spatial topology, and network distribution of the across-state distance (ASD). **c**, Average regional myelination (proxied by T1-weighted/T2-weighted image ratio) and the primary unimodal-to-transmodal FC gradient. Correlation to ASD assessed by spin permutation tests. **d**, Partial correlation between ASD, myelin, and the connectivity gradient. Hierarchical regression on the gradient loadings with myelin content and ASD as explanatory variables.

### Complexity dynamics are highly consistent in holdout data

Validation analyses in holdout data from the same participants showed that the distribution of signal complexity was remarkably consistent compared to the main analyses, yielding almost identical drop thresholds (Fig. 5a). Despite being highly dynamic metrics, the affinity of individual participants for complexity drops as well as their mean signal complexity were strongly correlated in main and holdout data (Fig. 5b). Moreover, independent clustering on the holdout data closely corroborated the distribution of neural complexity states, yielding a dominant high-complexity state and gradually more infrequent lower-complexity states (Fig. 5c), as in the main analyses. Furthermore, the degree to which complexity states were explored by participants (measured by the state dispersion index) as well as the spatial embedding of complexity states in the brain (measured by the across-state distance) were also highly consistent in holdout data (Fig. 5d).

**Fig. 5.**
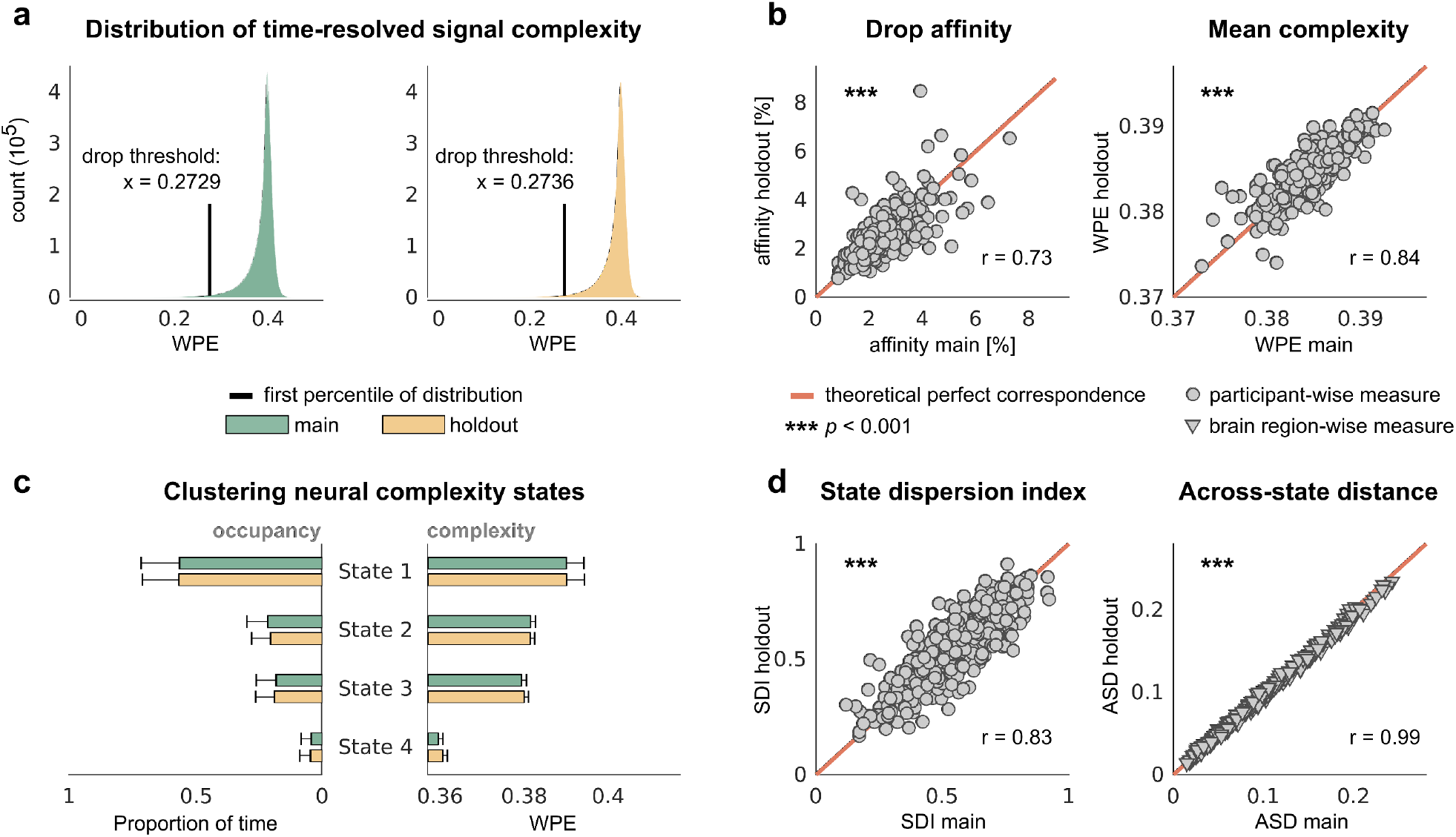
Validation of complexity measures in holdout data. **a**, distribution of time-resolved BOLD signal complexity in main and holdout data, resulting in nearly identical drop thresholds. **b**, consistency of participant-wise affinity for complexity drops and mean signal complexity (cf. Fig. 1b). **c**, state occupancy and state-wise signal complexity derived from independent clustering of neural complexity states in holdout data (cf. Fig. 3a). Error bars represent standard deviation. **d**, consistency of complexity state exploration as measured by the state dispersion index (SDI, cf. Fig. 3c) and the spatial embedding of complexity states as measured by the across-state distance (ASD, cf. Fig. 4b).

### Complexity-behavior associations

Lastly, we investigated the behavioral implications of the observed complexity dynamics, given that neural variability is increasingly recognized to carry important functional significance^25,39^. Based on the age-related reduction in drop affinity (Fig. 1b) and the link between complexity drops and functional integration (Fig. 3a,b), we expected lower complexity values to reflect better behavioral performance and lower age. To test this relationship, participant age and behavioral variables were subjected to a multivariate Partial Least Squares (PLS) analysis, including composite scores of crystallized and fluid abilities and the first principal components of cognitive, motor, and sensory task performance metrics (Extended Data Fig. 8). PLS returned a significant latent solution on the relationship between variance in complexity and differences in behavior (permuted p=2e-4). Supporting directional expectations, latent brain scores were positively related to latent behavioral scores (ρ=0.28), and in this latent space, neural complexity was positively linked to age and negatively associated with crystallized abilities, cognitive task performance and motor function (Fig. 6a). Notably, the associated bootstrap ratios showed that complexity-behavior associations were systematically constrained by functional networks (χ^2^(7)=95.4, p=9.5e-18), with strongest contributions by areas pertaining to the ventral attention and default mode network (Fig. 6b).

**Fig. 6.**
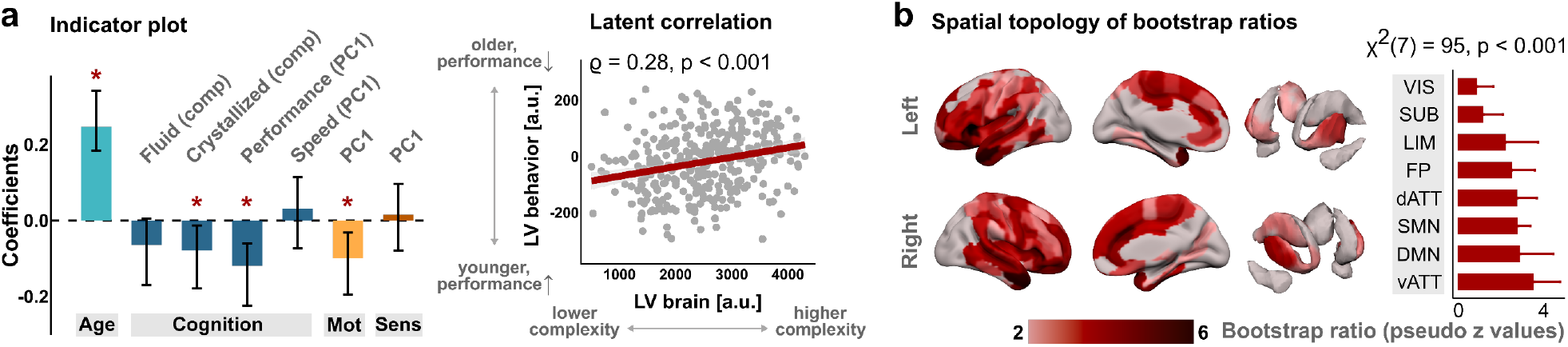
Neural complexity reflects inter-individual differences in age and behavior. **a**, Multivariate partial least squares correlation. Indicator plot (error bars: 95% CI) for age and behavioral variables (comp: composite score; PC1: first principal component; *: significant coefficient). Correlation of brain and behavior scores of the first latent variable (LV). **b**, Spatial topology, and network distribution of the corresponding bootstrap ratios.

## Discussion

Collectively, these findings delineate a unifying principle of brain organization, grounded in a spatiotemporal complexity architecture of neural activity. This human ‘complexome’ closes the gap between the variability of neural signals and several key properties of functional brain networks, with five immediate implications for our understanding of large-scale brain dynamics.

First, we show that resting-state brain activity is characterized by critical moments of neural regularity. Through time-resolved representation of regional BOLD dynamics, these episodes become visible as transient ‘complexity drops’ that are ubiquitously observed across scanning sessions, participants, and a variety of methodological parameters. Importantly, complexity drops occur spontaneously, are highly orchestrated over both time and space, and closely explain the coupling strength of functional connections as the degree to which brain regions engage in them simultaneously. Additionally, simultaneous drop engagement strongly differentiates between connections within, and connections across canonical resting-state networks, suggesting that functional subsystems are an expression of specific sets of regions dropping together particularly often. While these findings align well with the notion that functional connectivity is related to critical moments of neural activity^16,27,29^, they also characterize functional coupling as a graded, rather than an exclusively event-like process^40^. Although we find that regions show significant preference for dropping together with regions of their own canonical network, there is also substantial drop coincidence among regions that are traditionally assigned to distinct functional networks^12^, supporting the idea that brain regions can dynamically participate in different functional communities at different times^41,42^. In this view, our complexity framework may be able to bridge traditional ‘node-centric’ concepts of functional connectivity and more recent ‘edge-centric’ definitions of functional coupling^43^ because a high-amplitude co-fluctuation^27,29^ of two regional BOLD signals (edge level) here corresponds to those regions engaging in pattern regularity at the same time (node level).

Second, we show that complexity drops link two key phenomena of human brain activity: the dynamic propagation of neural patterns and the functional hierarchy of large-scale brain networks. While we observe that complexity drops can generally start anywhere in the brain, they consistently spread across the cortex along highly structured spatiotemporal propagation pathways. These propagation pathways intrinsically exhibit a principal hierarchy between unimodal systems that are central to the propagation, and transmodal systems that are less central to the propagation and tend to be their end points. Given that pattern propagation is thought to represent inter-regional communication^33,44^, our results suggest that such large-scale communication across brain networks may itself be hierarchical in nature, consistent with computational theories on information processing in the brain^45^. Furthermore, recent work has reported global arousal waves that similarly propagate from extrinsic, sensorimotor brain systems to intrinsic, higher-order systems, presumably reflecting spatiotemporal patterns of brain-wide excitability^28^. Although these reports were based on phase characteristics of the signal rather than amplitude information, the similarity of propagational pathways raises the possibility that moments of regularity could be linked to arousal-related states of excitability. Notably, the propagation of complexity drops also aligns well with the recently reported activity pulses in disused motor circuits^46^, which likewise occurred spontaneously and intrinsically spread through the motor network. Our results suggest that these pulses may represent an adaptive enhancement of a preexisting phenomenon – the dynamic propagation of regularity throughout the brain. Given our finding that these propagations inherently follow a principal functional hierarchy, such episodes may not be limited to impairment-induced plasticity^46^ but rather represent a more general mechanism by which the brain repeatedly and intrinsically self-maintains its functional architecture.

Third, we show that complexity drops define temporal states of neural activity, and that these neural complexity states dynamically modulate the coupling strength and topological configuration of the network in a moment-by-moment fashion. Given the unidirectionality of network construction – the correlation of neural signals is what defines the network in the first place –, our results suggest that these neural activity states may principally underlie the temporal states of the network. In this regard, we find that low-complexity states, in which many regions drop at the same time, yield functional states of high coupling, low modularity, and high efficiency in the network – consistent with previous findings on network efficiency dynamics^14^. In contrast, the default state of neural activity is a high-complexity state with precisely the inverse network configuration (i.e., low coupling, low efficiency, and high modularity). Notably, modularity and efficiency represent two complimentary network characteristics, where the former describes a segregated configuration thought to reduce biological cost and the latter implies an integrated configuration that enhances communication within the network^5^. The brain must balance both, however, yielding a cost-efficiency trade-off that is continuously re-negotiated^19^. Consequently, our findings indicate that the brain may implement such a trade-off with a default neural state of segregated activity that maintains cost-effectiveness, while the more infrequent low-complexity states ensure recurrent phases of functional integration.

Fourth, we show that the spatial embedding of complexity states comprehensively explains a well-established structure-function relationship in the brain^36^: the association between connectivity gradient loadings as a proxy of functional hierarchy^20,21^ and cortical myelination as a proxy of anatomical hierarchy^21,34,47^. Importantly, accounting for complexity states completely resolved this effect, whereas the complexity-myelin and complexity-gradient relationships persisted, showing that local anatomical properties and distributed functional properties of the network are linked through local functional dynamics.

Finally, we show that the observed complexity dynamics carry a number of inter-individual functional implications. On the one hand, we find several pronounced effects of participant age, including age-related reductions in (i) the affinity for complexity drops, (ii) the magnitude and diversity in how complexity drops are propagated throughout the brain, and (iii) the degree to which participants explore the space of neural complexity states. These findings suggest that neural dynamics grow increasingly more rigid with age, even over the narrow age range of early adulthood (22-35 years). On the other hand, these age effects are paralleled by an inverse association between brain signal complexity and indices of cognitive and motor performance, with strongest contributions by areas pertaining to the ventral attention network. Given that (i) an individual’s drop affinity strongly relates to overall signal complexity and age, (ii) complexity drops are tightly linked to phases of functional integration, and that (iii) the ventral attention network is thought to act as an intermediary system for switching between networks, these results suggest that a higher capacity for complexity drops may represent a beneficial aspect of brain functioning.

Overall, these findings lay out a coherent general framework to map regional neural dynamics to functional brain networks, with some key future questions readily following. First, the age effects observed in young adults raise the question of how complexity dynamics develop over the larger lifespan, including infancy, adolescence, and late adulthood. Similarly, the complexome provides a principled normative account to describe brain activity, with immediate opportunities to study clinical populations and altered mental states. Moreover, the present study focuses on resting-state brain activity as assessed by fMRI, warranting further study on how complexity dynamics change during cognitive task engagement, and what the correlates of complexity drops are in electrophysiological recordings with higher temporal resolution. Finally, complexity dynamics are remarkably stable within participants, raising the question of how these dynamics relate to individual functional ‘fingerprints’^48,49^ and precision mapping of individual brain organization^50^.

In sum, our study speaks to a model of the brain in which its intricate functional architecture is tightly linked to moments of neural regularity, with several new avenues to understand large-scale brain dynamics.

## Online Methods

### Data and preprocessing

Data were obtained from the Human Connectome Project (HCP)^51,52^, including minimally processed resting-state fMRI^52,53^, diffusion-weighted images (DWI), T1-/T2 image ratios^47^ as well as demographics and behavioral scores from 343 participants (205 females, 138 males). MRI data were acquired in a 3T scanner at Washington University in St. Louis with multiband echo-planar imaging (1200 volumes, TR = 0.72s, 2mm isotropic voxels). Two runs of approximately 15 minutes each, one with right-to-left (RL) and one with left-to-right (LR) phase encoding protocol, were acquired on two consecutive days, resulting in a total of four resting-state datasets per participant. Spatial distortion correction was applied as provided in the HCP.^52^ Of the four scans, one encoding direction was chosen at random for day 1 and 2, respectively, resulting in two scans per participant with balanced phase encoding distribution for the main analyses (day1-LR: n=168, day1-RL: n=173, day2-LR: n=166, day2-RL: n=177; Pearson’s χ^2^-test: χ^2^=0.05, p=0.82), with the remaining two scans serving as a holdout dataset for the validation analyses presented in Fig. 5. Two scans from different participants were excluded because of incomplete data, resulting in a total of 684 scans for the main analyses. All acquisition parameters and processing pipelines for these data are described in detail elsewhere^52–56^. Bandpass filtering [0.01 Hz to 0.1 Hz] and z-scoring were applied to each voxel before timeseries extraction. The effect of filter settings was investigated in Extended Data Fig. 3. Regional BOLD timeseries were extracted with the Brainnetome (BNA) atlas^57^, which includes 246 cortical and subcortical regions of interest (ROI). Assignment of ROIs to seven canonical resting-state networks (visual (VIS), somatomotor (SMN), dorsal attention (dATT), ventral attention (vATT), default mode (DMN), frontoparietal (FP), and limbic (LIM)) is provided with the BNA template^57^. This mapping is derived from the cortical network parcellation by Yeo et al.^12^ and is available from www.brainnetome.org/resource. Regions 165, 177, and 178 (located in the left insular gyrus, left cingulate, and right cingulate, respectively) are not labelled in the template and were manually assigned to the frontoparietal network based on their overlap with the Yeo parcellation. Furthermore, regions in the basal ganglia, thalamus, hippocampus, and amygdala were subsumed as subcortical parcels. Finally, we investigated the sensitivity of the observed complexity dynamics to methodological choices of functional parcellation and timeseries extraction (volumetric vs. surface-based extraction of cortical timeseries). To this end, we compared the BNA parcellation to two volumetric atlases of higher and lower spatial granularity (after Shen et al.^58^ with 268 ROIs, and after Shirer et al.^59^ with 90 ROIs) as well as to the surfaced-based MMP atlas with 360 cortical ROIs after Glasser et al.^60^, yielding highly convergent results (Extended Data Fig. 4-5).

### Time-resolved estimation of signal complexity

Signal complexity of BOLD activity was calculated as weighted permutation entropy (WPE) through symbolic encoding of the timeseries vectors^30^. WPE is an amplitude-sensitive extension of permutation entropy (PE)^61^, an information-theoretic quantity that captures the degree of pattern irregularity as the Shannon entropy^62^ on the occurrence of symbolic motifs within a timeseries.

WPE is defined as

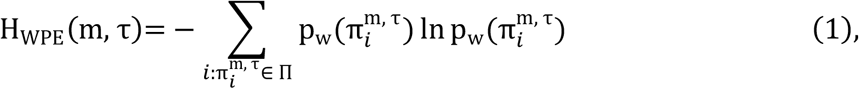

where m is the length of symbolic motifs in the timeseries, *τ* is a lag parameter indicating the number of time points to shift along the timeseries, 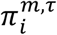 represents the *i*’th symbolic motif out of the set of possible motifs Π given the motif length, and 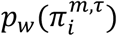 is the variance-weighted relative frequency of motif *i*, as detailed by Fadlallah et al.^30^. Following methodological considerations^30,63^ and previous applications of WPE to neural data analysis^64,65^, a motif length of m = 3 and a lag parameter of *τ* = 1 was applied to compute the motif distribution over the signal vector of interest, and WPE bit values were normalized to lie in [0, 1]^30^. Notably, by leveraging both pattern diversity (assessed by PE) and amplitude information (assessed by the standard deviation over the BOLD vector), WPE captures signal dynamics that remain undetected when considering only one of these characteristics (Extended Data Fig. 1-2). A further advantage of WPE lies in its ability to accommodate a time-resolved approach, based on the comparatively low number of timeseries samples needed to achieve stable estimation^63,66^. Accordingly, we here applied a sliding window approach, where the entire BOLD timeseries in divided into temporally contiguous windows of a prespecified length and overlap. In the main text, we report the findings for a window length of 60 TRs (43.2 seconds) with 95% overlap, yielding a window-to-window temporal resolution of 3 TRs (2.16 seconds) and a time-resolved signal complexity vector of 380 windows. To investigate the sensitivity of the observed complexity dynamics to the windowing parameters, we analyzed window lengths of 60, 90, and 120 samples with 95% and 90% window overlap, respectively, yielding highly convergent results (Extended Data Fig. 4).

Based on the ubiquitously observed pattern of predominant high complexity with recurrent complexity drops, we defined a threshold of drop engagement as the first percentile of the total WPE distribution (critical WPE = 0.273; cf. Fig. 5). Accordingly, whenever a region’s signal complexity met this value for a given BOLD window, it was counted as engaged in a complexity drop in that window (Fig. 1a). Notably, whether this thresholding procedure was applied across all participants or as participant-specific percentile thresholds made essentially no difference to the spatial distribution of where complexity drops occurred (r = 0.99, p<0.001). We then investigated to which extent brain regions coincide in their drop engagement over time. To this end, we iterated over all BOLD windows in the data set and counted those regions that met the drop threshold simultaneously in each window. This count matrix was normalized by the maximum coincidence count, yielding drop coincidence values from 0 to 1. The pairwise drop coincidences between any two regions were then subjected to correlation analyses with the corresponding functional connectivity values (Fig. 1c). To investigate how the likelihood to engage in complexity drops is distributed over brain regions and age groups, we furthermore calculated time-resolved drop affinity for each scan. To this end, a binary matrix (BOLD windows over ROIs) was created, containing ones whenever a region’s signal complexity met the drop threshold in a particular window, and zero otherwise. These affinity matrices were then averaged to estimate participant-wise and region-wise drop affinity. Note that the time-resolved affinity vector for individual scans also served the observation of brain-wide engagement in complexity drops (Fig. 2a) and the subsequent propagation analysis.

We furthermore assessed the degree of similarity between a region’s complexity timeseries with its contralateral equivalent (Fig. 1d). To this end, we defined an interhemispheric symmetry index (ISI), which is computed as the correlation coefficient of a region with its contralateral equivalent, weighted by the proportion of the 246 BNA regions that were less strongly correlated with that region. Accordingly, a region with high cross-hemispheric similarity shows many regions that are less correlated with it than the contralateral equivalent, yielding a weighting factor close to one. In contrast, if there are many regions that exhibit a higher correlation to the target region than the contralateral ROI, this weighting factor decreases, resulting in lower ISI values.

### Descriptive statistics

Group-level comparisons (e.g., by age or by resting-state networks) were carried out with the nonparametric Kruskal-Wallis test^67^, which rests on observation ranks and applies to n>2 groups. Effects were considered statistically significant at a level of α=0.05 for all tests. Pairwise comparisons were conducted with rank sum tests^68^, adjusting for multiple comparisons using the false discovery rate (FDR). Associations between continuous variables were assessed with parametric or nonparametric correlation tests, depending on the underlying distributions. Participant age was given in the age groups of 22-25, 26-30, 31-35, and >35 years; for age-wise comparisons, however, participants over 35 years were excluded, as this only applied to n=3 individuals. For all assessments of directional effects, two-tailed tests were applied.

### Propagation analysis

The propagation of complexity drops across the brain was formalized as a graph theoretical problem. Here, each individual propagation was modelled as a directed graph where nodes represent brain regions engaged in complexity drops and directed edges represent the progression in time from one BOLD window to the next.

To identify drop cascades in the dataset, a stepwise search procedure was applied. First, all BOLD windows with at least n=10 regions simultaneously engaged in complexity drops were identified in the time-resolved affinity vector. If there were several temporally contiguous windows that met this criterion, the window with the maximum number of engaged regions was defined as the peak layer in the directed graph. From this peak layer, a window-by-window backward and forward search identified the neighboring windows in which the number of engaged regions increased until reaching the peak layer (propagation phase) and decreased after the peak (fade phase), respectively. The minimum cascade length thus included three contiguous windows (initialization – peak – fade). Although very rare, instances in which the windows directly before or after the peak showed no engaged regions were discarded. To investigate the spread of complexity drops in the propagation phase, individual directed graphs were constructed from the initialization window – in which the engaged regions represent the graph’s source nodes – to the peak window (Fig. 2b).

For all windows from initialization to peak, the newly engaging regions from one window to the next were registered to compute the empirical transition probability, where all newly engaged regions in window *i* obtain directed edges to all newly engaging regions in window *i* + 1. The region-by-region transition probability matrix (TPM) was then constructed based on the path-weighted edges in the propagation graph. Here, edges in temporally contiguous windows were assigned a weight of 1, while connections in non-neighboring windows were assigned the inverse of the path length (e.g., ½for connections from window *i* to window *i* + 2) to account for the temporal evolution of the spread. For instance, if engagement of a region A is frequently followed by engagement of region B, but through variable intermediate engagement of regions C, D, etc., this is missed in a binary TPM, as only neighboring window are considered. Construction of propagation graphs and TPMs over individual cascades subsequently allowed for participant-wise and group-level analyses: to capture cascade origination, the source node probability of each brain region was computed as the rate of occurrence in the initialization window. The diversity of source nodes was calculated as the percentage of unique brain regions ever initializing a participant’s cascades. The average TPM over individual spreads yielded the directed group-level propagation network in Fig. 2e, representing the 95^th^ percentile of path-weighted transition probabilities. Graph construction and topological analyses on the propagation network was implemented with the igraph package (version 1.2.5) for R^69^. The hub structure of the network was quantified as Kleinberg centrality, an extension of eigenvector centrality for directed networks^70^. Geodesic distances were computed for every node-to-node comparison in the propagation network and averaged for all combinations within and across resting-state networks. The average distance matrix was then subjected to hierarchical clustering using the default complete linkage method^71^. As this approach revealed a highly suggestive order from unimodal to transmodal networks, we explicitly investigated this cluster structure by defining a unimodal and a transmodal cluster group, subsequently tested with a Monte Carlo simulation using the sigclust package^72^. The null hypothesis of this cluster test is that the data emanate from a single Gaussian distribution. To assess this hypothesis, 5000 Gaussian samples were created to estimate the distribution of the cluster index^72^, a test statistic defined as the sum of within-class sums of squares about the mean in relation to the total sum of squares about the overall mean (cf., https://rdrr.io/cran/sigclust/man).

### Clustering of complexity states

The time-resolved signal complexity matrix (rows: 684 scans x 380 windows = 259920 observations; columns: 246 ROIs) was subjected to unsupervised structure detection to investigate temporally discrete complexity states. To this end, we applied the MATLAB-inbuilt k-means clustering algorithm with a maximum of 1000 iterations, 20 replicates with random initial positions to avoid local minima, the k-means ++ heuristic for centroid initialization, and the squared Euclidean distance as the target metric to be optimized. Based on the observed complexity dynamics (Fig. 1a; online repository: https://osf.io/mr8f7), we expected a predominant high-complexity state, an infrequent low-complexity state, and a varying number of intermediate complexity states. As common cluster evaluation indices to determine the optimal k of expected clusters resulted in heterogeneous estimates between k=3 and k=5 depending on windowing parameters and the optimization criterion applied, we specified k=4 (i.e., two intermediate complexity states) and ran comprehensive validation analyses for k=3 (i.e., one intermediate state) and k=5 (i.e., three intermediate states). Results reported in the main text were extremely robust against the choice of k, including the occupancy-complexity relationship, the impact of complexity states on network strength and topology, state dispersion, as well as the spatial topology of complexity states (whose spatial heterogeneity was assessed based on the modified likelihood ratio test^73^, MLRT) and its link to myelination and the primary functional connectivity gradient (Extended Data Fig. 6-7).

#### State dispersion index (SDI)

To assess the degree to which the empirical state visits of an individual participant were distributed across the *k* possible complexity states, we defined a state dispersion index (SDI), calculated as

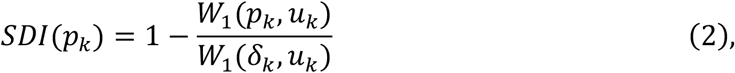

where *W*_2_(*p*_1_, *u*_1_) corresponds to the first Wasserstein distance^74^ between the empirically observed discrete distribution of state occupancy *p*_1_ (Fig. 3a-c) and the theoretical maximum dispersion distribution *u*_1_ = *U*(1, *k*) (i.e., the uniform distribution in which each state is visited with a frequency of 1/*k*). This distance is normalized by *W*_2_(δ_1_, *u*_1_), which is given by the Wasserstein distance between the theoretical minimum dispersion distribution δ_1_ (i.e., a zero-entropy degenerate distribution in which only one of *k* states is ever visited) and the uniform distribution. Consequently, the *SDI* lies in [0,1], and is bounded by 0 if only one state was ever visited by the participant (minimum state exploration), and 1 if the empirical state occupancy is identical to the uniform (maximum state exploration).

#### Across-state distance (ASD)

To characterize the spatial topology across the estimated complexity states, we defined an index of across-state distance (ASD). Let *K* denote the set of unique centroid pairs (*i, j*) in *k*-dimensional state space. For each brain region, the ASD is computed as the cumulative Euclidean distance *D* over all pairs of centroids *C* in *K*:

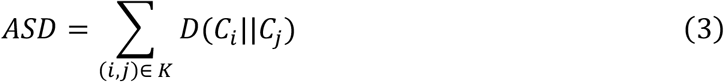

For the Brainnetome parcellation, this yields a 1 × 246 vector, where higher values indicate brain regions that show greater centroid-to-centroid distances and thus more pronounced differences in signal complexity over the estimated complexity states. Note that the resulting ASD topology was very robust against the choice of k (Extended Data Fig. 6-7), and that this vector constituted the input for the subsequent correlation analyses linking state topology to cortical myelination and the primary functional connectivity gradient.

### Functional connectivity and network topology

Static functional connectivity was estimated as the ROI-by-ROI Pearson correlation matrix over the entire resting-state recording. These matrices were averaged over runs and participants to obtain the group-level functional connectivity matrix for investigating the association to drop coincidence and for the construction of the macroscale connectivity gradient. Furthermore, dynamic functional connectivity was computed as the correlation matrix over each BOLD window, resulting in a 246×246×380 array for the specified window parameters. Dynamic connectivity strength was calculated as the mean over these window-wise matrices, yielding a 1×380 vector. The topology of the time-resolved functional networks was assessed with the Brain Connectivity Toolbox^75^, available from www.brain-connectivity-toolbox.net. Network modularity as a measure of functional segregation was estimated through Louvain community detection (community_louvain.m) with asymmetric treatment of negative weights as previously recommended^75^ and with a γ-parameter of 1.05 to accommodate the ability to detect smaller modules^75^. Furthermore, global efficiency as an estimate of functional integration was computed using the efficiency_wei.m function, with negative weights discarded^75^. This approach yielded vectors of time-resolved modularity and global efficiency (1×380 BOLD windows) for each resting-state recording. These time-resolved vectors constituted the input for the calculation of both the participant averages (Fig. 3a) and the window-to-window transitions (Fig. 3b) of these network measures, where the corresponding complexity state vector from the clustering output served as a state-wise mask.

### Structural connectivity estimation

Structural connectivity matrices were computed using probabilistic tractography as implemented in the FMRIB’s Diffusion Toolbox ProbtrackX GPU program^76^. Diffusion-weighted data provided as the Diffusion BedpostX package were available for 340 of the 343 participants in the study population. These data were preprocessed as previously described^52^, including registration to native space, movement and eddy currents correction, and application of BedpostX to model white matter fiber orientations for probabilistic tractography. The connectivity distribution is then computed using iterations of streamlines propagated from seed regions to target regions. Briefly, a single propagation entails moving along a streamline by steps of a specified length, evaluating for exclusion or termination criteria at each step, and continuing this process until criteria for exclusion (streamline discarded) or termination (propagation ended but streamline retained) are met. The output is a matrix which quantifies the number of non-discarded streamlines between a seed and target.

In line with the functional analyses, BNA regions were specified as seed ROIs to guide tractography. Network mode was applied in ProbtrackX to compute a ROI-by-ROI connectivity matrix, where rows represent seed ROIs and columns correspond to target ROIs. Each BNA region was input as a binary mask in MNI space. Since the seed ROIs (standard) and processed diffusion data (native) were not registered to the same space, bidirectional participant-specific nonlinear transformations between standard and structural space as provided by the HCP were passed to ProbtrackX. Following the recommendation that the inclusion of additional anatomical priors increases the biological plausibility of modelled white matter tracts^77^, we included participant-specific gray matter (GM) masks from Freesurfer as termination masks (--stop). By terminating a propagated streamline as soon as it enters this GM mask, streamlines are forced to terminate near the boundary between GM and WM, instead of travelling further into a GM region. Streamline termination parameters were applied as ProbtrackX defaults: streamline curvature threshold exceeded (0.2, ∼80 degrees), streamline path returns to a point which it already intersected previously (--loopcheck), streamline exits the nodiff_brain_mask (BedpostX output), and maximum number of steps per streamline (2000) reached using a step length of 0.5 mm.

For each voxel in a seed region, 1000 samples were propagated. Distance correction (--pd) was applied to compensate for the bias that the probability of a streamline successfully reaching the target ROI decreases as the distance between a seed and target ROI increases. Distance correction as implemented in ProbtrackX adjusts the connectivity distribution between a seed-target ROI pair by multiplying the number of successful streamlines between the ROIs by the average path length of the streamlines. Finally, ROI-by-ROI connectivity matrices were then normalized using the probabilistic streamline connectivity index (PSCI) procedure, as previously described^78^. The PSCI scales the streamline counts between ROI pairs based on the number of propagated and successful streamlines as well as the size of both the seed and target ROIs.

### Myelination estimation

Cortical grayordinate myelin maps obtained from the T1w/T2w image ratios^47^ were used to calculate the average myelin content of ROIs in the atlas template. Since our parcellation approach did not benefit from the higher resolution of the 164k images, we analyzed the unsmoothed, bias-corrected 32k areal-feature-based aligned images (‘MyelinMap_BC_MSMAll.32k_fs_LR.dscalar.nii’) using the Multimodal Surface Matching algorithm^79^. Parcel-wise myelin content was obtained using the Connectome Workbench (command: -cifti-parcellate) by averaging over the myelin map within the respective BNA regions.

### Gradient construction, spin permutation, and hierarchical regression

The macroscale connectivity gradient was computed on the group-average functional connectivity matrix. As only the cortex is mapped to the FreeSurfer sphere for later spin permutation, the input matrix was restricted to the 210×210 cortical ROIs of the BNA. Gradient analysis was implemented with the BrainSpace toolbox^80^ for neuroimaging and connectomics datasets. Following reference^20^, cosine similarity was applied to compute the affinity matrix. A total of 10 components, diffusion embedding for non-linear dimensionality reduction (diffusion time of 0, alpha parameter of 0.5), and 90% region-wise feature sparsity were used for gradient construction, following the default recommendations^80^. The gradient fit on these data is displayed in Fig. 4c of the main text and precisely identified the principal unimodal-to-transmodal connectivity gradient first reported by Margulies and colleagues^20^. To examine the association between these gradient loadings and the other cortical features (myelination, across-state distance), we applied spin permutation correlation of the respective cortical maps. This approach consists in a series of random spherical rotations that preserve the spatial autocorrelation of the data, resulting in empirical null models that counteract the potential inflation of statistical significance in simple univariate tests^80–82^. Here we implemented a variant of this procedure that specifically applies to parcellated instead of vertex-wise cortical maps^83^. Nonparametric correlations over 5000 random rotations were computed to estimate the empirical distribution of the test statistic for the three comparisons between gradient loadings, cortical myelination, and the across-state distance.

The empirically observed associations across these cortical features were furthermore subjected to a nonparametric partial correlation analysis using the ppcor package for R^84^ (Fig. 4d). As this approach suggested high explanatory power of complexity states, these analyses were corroborated with a hierarchical regression approach, where variance in gradient loadings was modelled as a function of cortical myelination (compact model 1), across-state-distance (compact model 2), or both (augmented model 3). These individual linear models were then compared through F-tests for nested models with the lmSupport package for R (https://cran.r-project.org/web/packages/lmSupport/lmSupport.pdf).

### Complexity-behavior associations

To assess associations between signal complexity and participant characteristics, average WPE values per ROI and participant were tested against age and indices of cognitive performance, motor skills as well as sensory scores, as provided with the HCP data. To ensure data quality and minimize the impact of non-normally distributed variables, we transformed response time data to speed via inversion, subjected skewed data to log transformations (if absolute skewness >1), excluded data from participants with missing values in at least one variable, and removed outliers (mean ± 4 standard deviations). In total, data from 13 participants were thus excluded, leaving n=330 participants for behavioral analyses. To this end, we tested the multivariate relationship between BOLD signal complexity and individual participant measures by means of a partial least squares (PLS) analysis^85–87^ (Fig. 6). We applied a behavioral PLS approach that allows for the estimation of multivariate correlations between a three-dimensional brain variable (average WPE per ROI and participant) and multiple behavioral measures (Extended Data Fig. 8). To focus this analysis on general features of behavioral and cognitive abilities, we performed principal component analyses (PCA) on the respective sets of single measures of performance and speed in cognitive tasks, motor performance, and sensory skills. For each of the four domains, the first principal component accounted for more than 35% of total variance, and component loadings were strictly positive, resulting in four principal component scores per participant that entered PLS analyses alongside individual age as well as composite scores of fluid and crystalized cognition, as provided with the HCP data, such that seven behavioral variables were used in the PLS analysis.

In brief, PLS works via the calculation of a correlation matrix that captures the between-participant correlation of the target brain measure in each region and the behavioral metrics of interest (matrix size: N_regions_ x N_behavior_ = 246 × 7). Next, this rank-correlation matrix is decomposed using singular value decomposition (SVD), resulting in N_behavior_ x N_behavior_ latent variables. This approach produces two main outputs: (i) a singular value for every latent variable, indicating the proportion of cross-block variance explained by the latent variable, and (ii) a pattern of weights (N_regions_) representing the rank correlation strength between WPE and behavioral measures. The multiplication (dot product) of these weights with region-wise WPE yields brain scores reflecting the between-participant correlation of complexity and behavioral metrics. Statistical significance of these brain scores and underlying latent variables was tested by permuting behavioral measures across participants and recalculating the singular value of each latent variable (5000 permutations). To furthermore estimate the robustness of the calculated weights, a bootstrap procedure was applied (5000 bootstraps with replacement). The division of the empirical weights by the bootstrapped standard error yields bootstrap ratios (BSR). These BSR values estimate the robustness of observed effects on a region-wise level and can be interpreted as values from a z-distribution. Hence, BSR values exceeding 1.96 relate to a correlation between the latent brain scores (weighted average WPE) and the latent behavioral scores (weighted behavioral metrics) at a p-value of p<0.05, and likewise p<0.01 for BSR values exceeding 2.7. Finally, bootstrap resampling was also applied to estimate the 95% confidence intervals for the observed indicator correlations between WPE and behavioral measures.

### Data and code availability

All data analyzed here are publicly available from the Human Connectome Project (www.humanconnectomeproject.org). Analysis code is available from the corresponding authors and will be made public upon acceptance of the manuscript. Further information including additional validation analyses are available from the online repository https://osf.io/mr8f7.

## Acknowledgments

We thank Maron Mantwill for assistance in data management. This work was supported by the German Ministry of Education and Research (BMBF) grant 13GW0206D (SK), the Cusanuswerk foundation (NS), the Berlin School of Mind and Brain (AR), the German Research Foundation grant IRTG2150 (MG), the Max Planck UCL Centre for Computational Psychiatry and Ageing Research (LW, DG), the German Research Foundation Emmy Noether Programme (DG) as well as German Research Foundation grants 327654276 (SFB 1315), FI 2309/1-1 (Heisenberg Programme), and FI 2309/2-1 (CF).

## Author contributions

SK, NS, and CF conceived the study. SK, NS, AR, and MG processed imaging data. LW provided methodological expertise on estimation of weighted permutation entropy. SK, NS, LW, AR, and MG analyzed the data. SK designed data analysis and developed novel analysis methods (propagation analyses, state exploration, spatial embedding of neural states) with feedback from NS, LW, DDG, and CF. NS, AR, and LW validated data analysis. LW and DDG contributed methodology and implemented the PLS analysis. SK, NS, and AR visualized results. All authors interpreted results. SK drafted the manuscript with contributions from NS, LW, AR, and MG. DDG and CF supervised the study. All authors revised, edited, and approved the manuscript.

## Additional information

Extended Data Figures 1-8 Supplementary Movie 1

**Extended Data Fig. 1.**
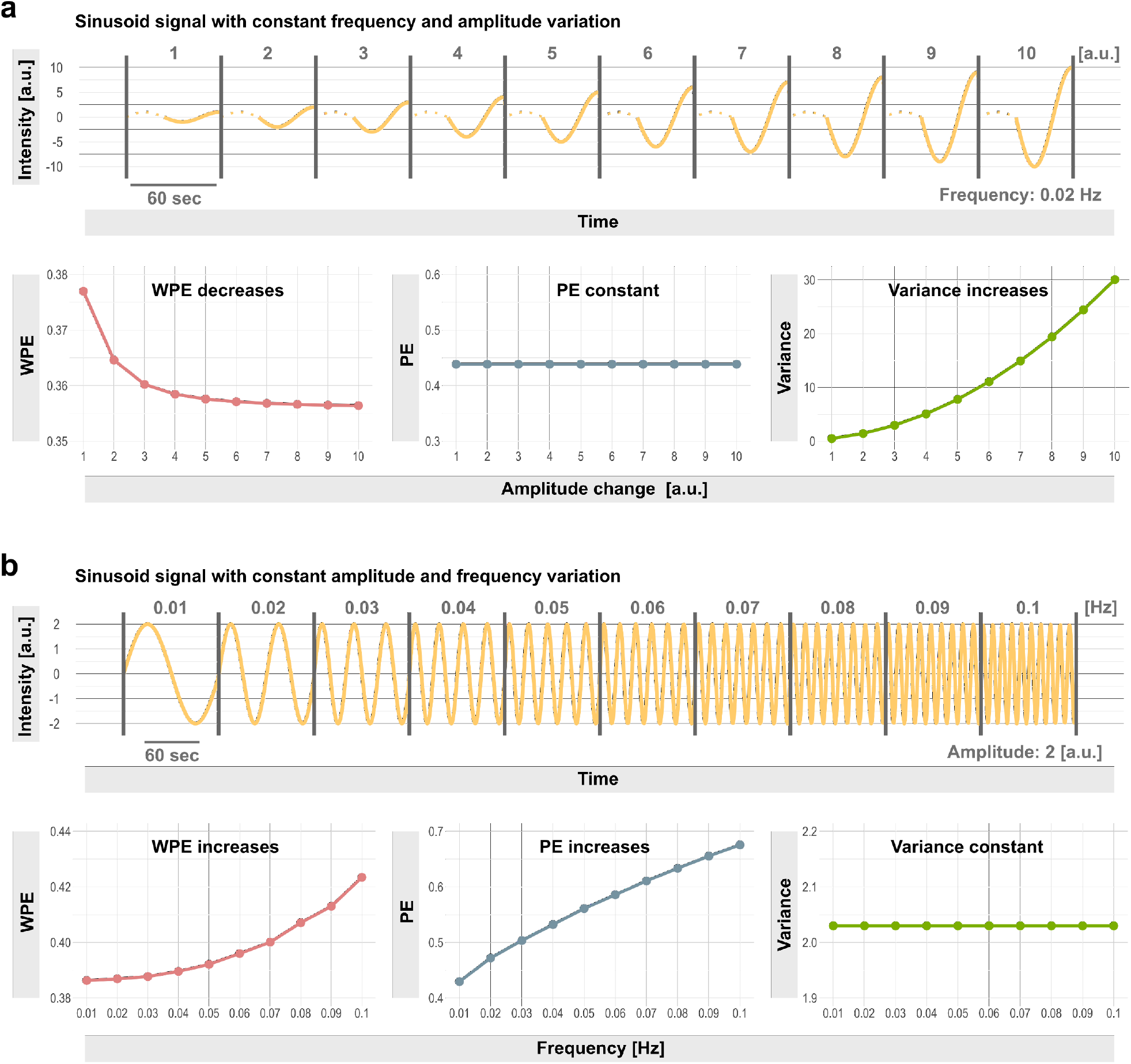
Weighted permutation entropy reflects amplitude and frequency variation that remains undetected when considering only pattern diversity or signal variance alone. **a**, Simulated sinusoid signal with constant frequency in the range of real BOLD signals (0.02 Hz) and with systematically increasing amplitude variation that results in changes of weighted permutation entropy (WPE) and amplitude variance, whereas pattern diversity alone (permutation entropy, PE) remains identical. **b**, Simulated sinusoid signal with constant amplitude and systematically increasing frequency variation that results in changes of weighted permutation entropy (WPE) and pattern diversity (PE), while signal variance remains unchanged. Note that WPE values monotonically increase with the higher frequency content, reaching about 0.42 normalized bits at 0.1 Hz (identical to the applied low-pass filter settings in BOLD preprocessing), consistent with the empirically observed upper limit of the complexity timeseries derived from real BOLD signals (Figure 1a; online repository: https://osf.io/mr8f7; Extended Data Figure 3-4).

**Extended Data Fig. 2.**
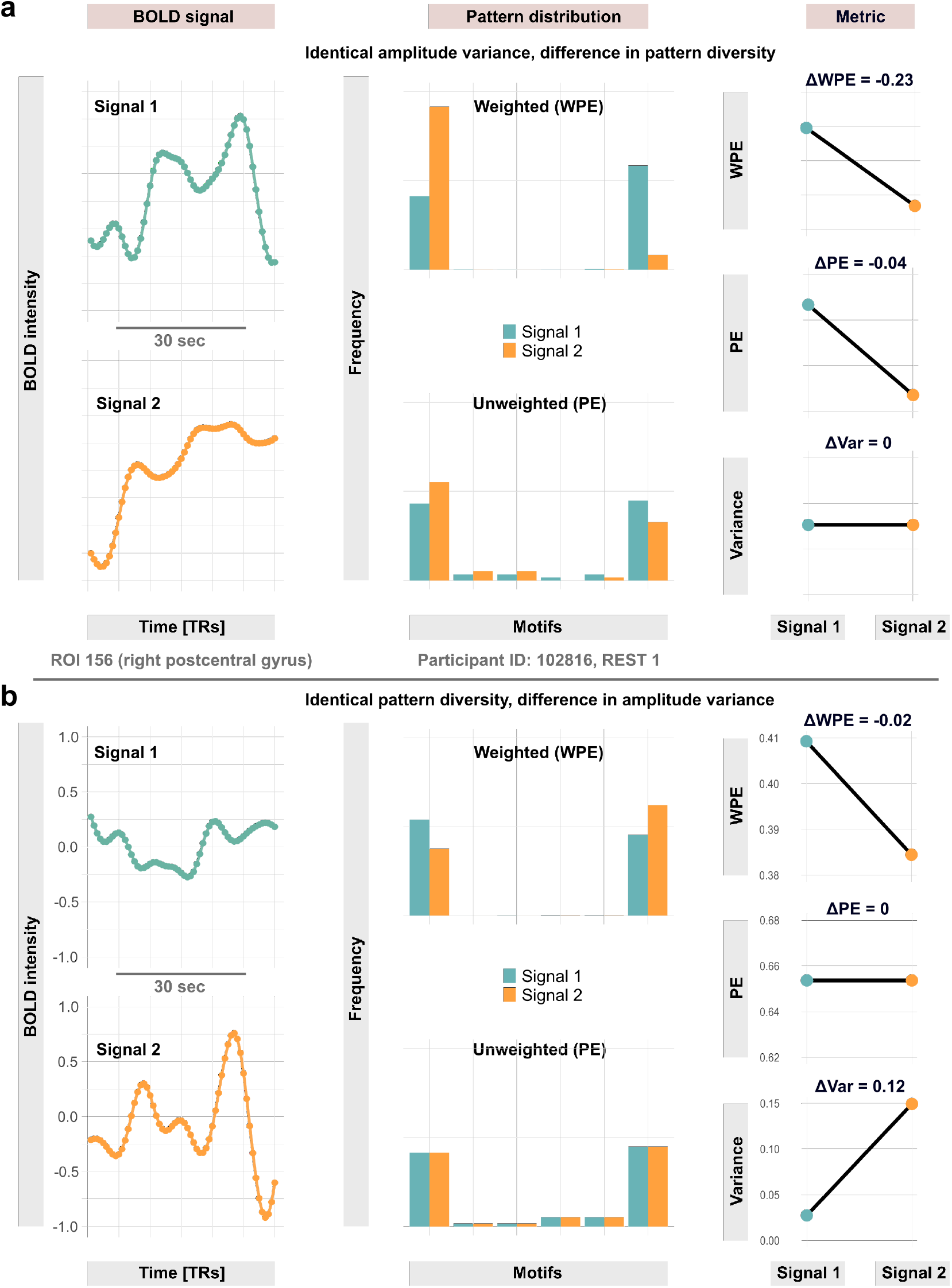
Weighted permutation entropy tracks BOLD signal dynamics that remain undetected when considering only pattern diversity or signal variance alone. **a**, BOLD signals with identical amplitude variance but varying pattern diversity (permutation entropy, PE). **b**, BOLD signals with identical pattern diversity but varying amplitude variance. All signal snippets correspond to non-overlapping BOLD windows from the right postcentral gyrus of a representative resting-state recording. Columns show BOLD signals (left), the associated variance-weighted pattern distributions (middle), and the change in signal metrics (right).

**Extended Data Fig. 3.**
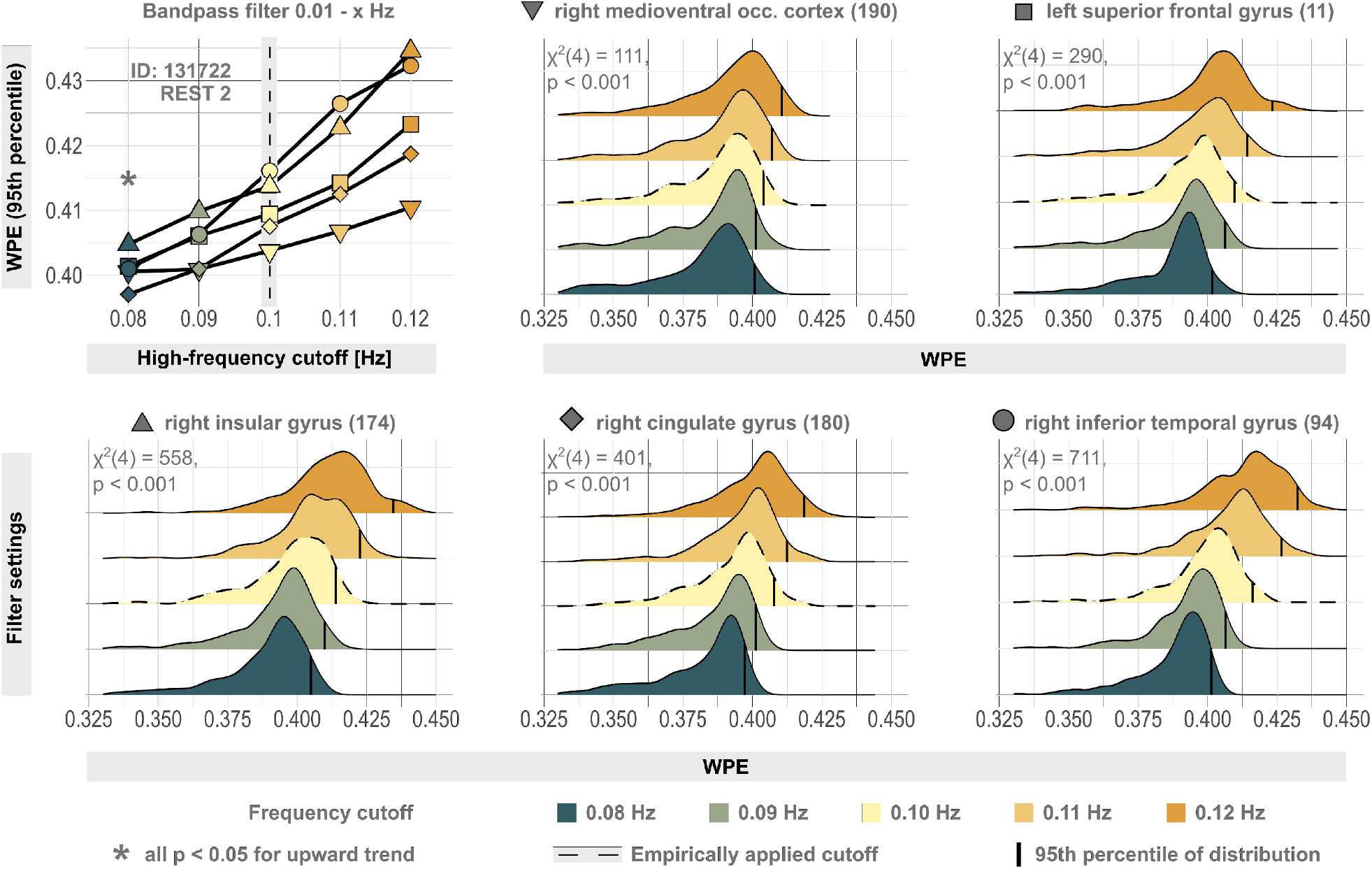
Upper-frequency content scales the distribution of weighted permutation entropy. The upper left panel shows the 95th percentile of weighted permutation entropy (WPE) values from five randomly selected ROIs over different bandpass filter settings in a representative resting-state dataset. Brainnetome IDs of regions are given in brackets. In line with the frequency-sensitivity in pure sine signals (Extended Data Figure 1), systematic variation of the high-frequency cutoff gradually scales the upper limits of WPE values, which becomes apparent as significant monotonic upward trend in nonparametric Mann-Kendall tests and a systematic right-shift of the time-resolved WPE distributions for individual regions. Note, however, that the heavy left tails (corresponding to complexity drops, cf. Figure 5a) are unaffected by scaling of the distribution. For the applied low-pass cutoff of 0.1 Hz in the main text (dashed lines), maximum WPE values converge around 0.4 both in simulated and real signals, in line with the empirically observed upper limit of individual complexity timeseries (Figure 1a and 5a; online repository: https://osf.io/mr8f7; Extended Data Figure 4).

**Extended Data Fig. 4.**
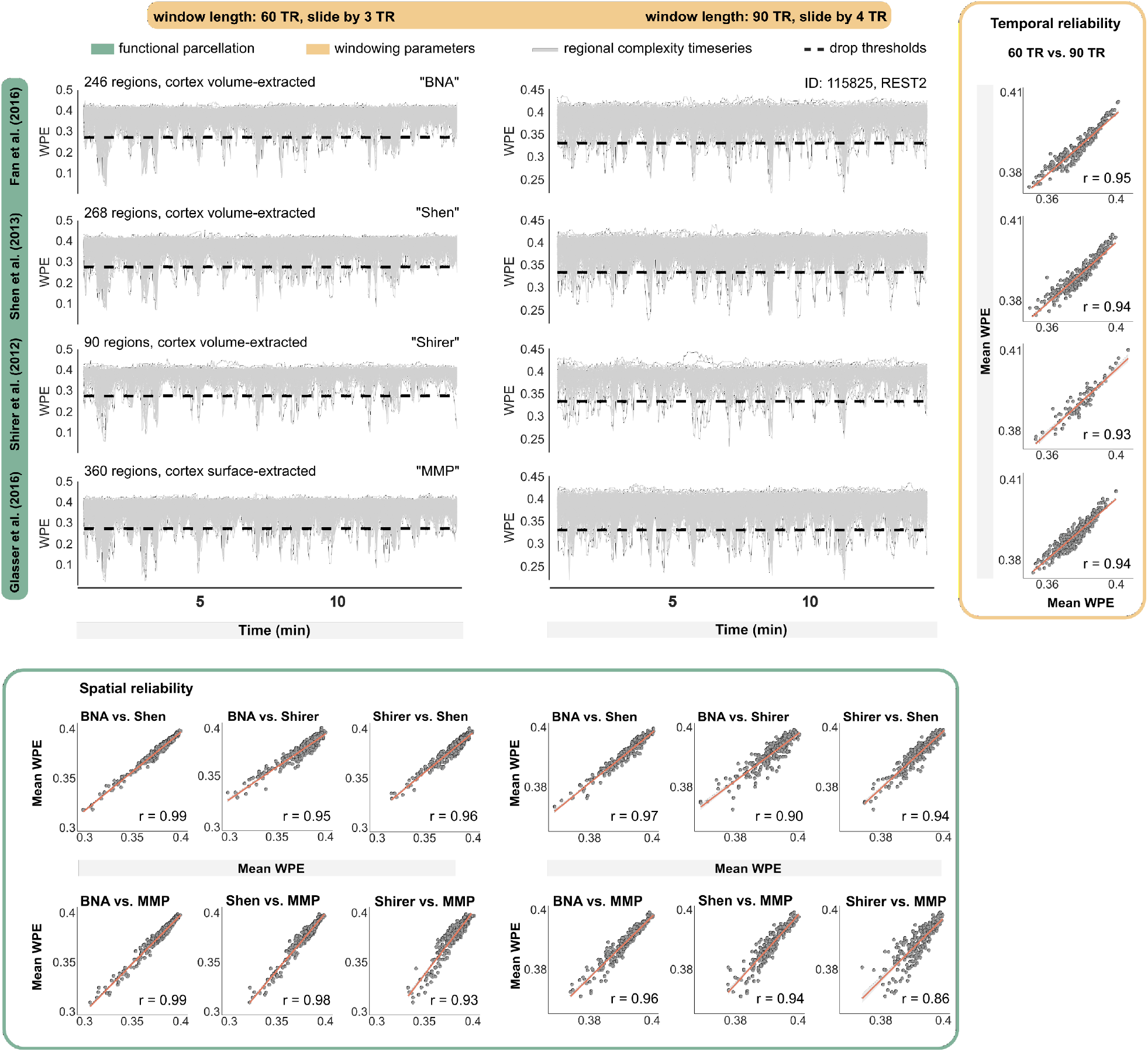
Resting-state complexity dynamics are robust across functional parcellation granularity, windowing parameters, and timeseries extraction methods. Time-resolved weighted permutation entropy (WPE) from an exemplary HCP scan. The data were parcellated with three common volumetric brain atlases of varying spatial granularity (Brainnetome after Fan et al.^57^, Shen et al.^58^, and Shirer et al.^59^) as well as one atlas in which subcortical timeseries are extracted volumetrically but extraction of cortical signals is surface-based (Glasser et al.^60^). Furthermore, different windowing parameters for computing WPE timeseries are applied (60 TRs sliding by 3 TRs, and 90 TRs sliding by 4TRs). Spatial reliability is calculated as the correlation over time points for every combination of functional atlases, and temporal reliability is computed as the correlation of mean WPE values of every ROI for the two windowing parameters. Results in the main text are based on the Brainnetome atlas and 60 TR windows sliding to the next window by 3 TRs (upper left panel here).

**Extended Data Fig. 5.**
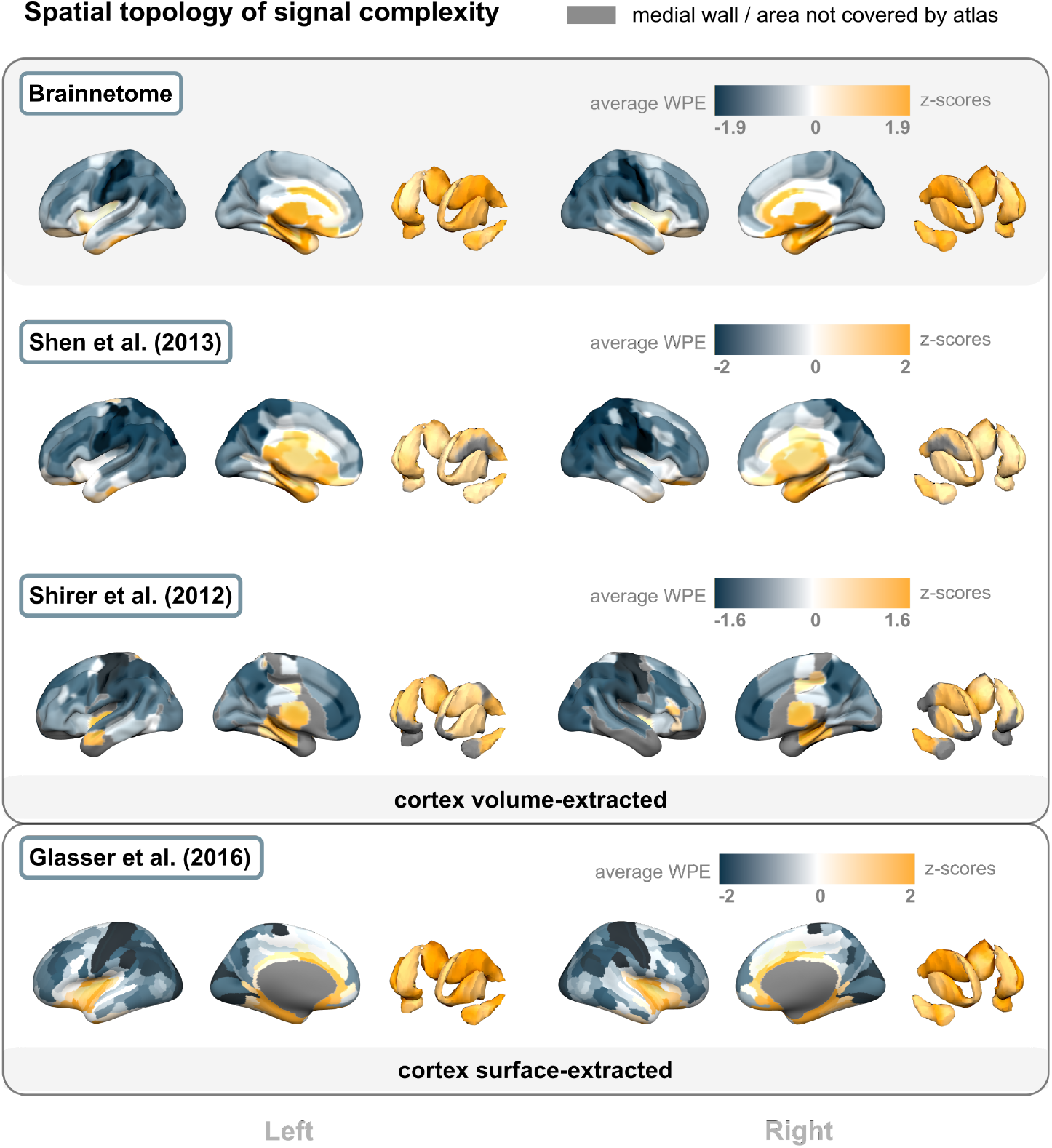
The spatial topology of signal complexity is robust across functional parcellation granularity and timeseries extraction methods. Spatial distribution of weighted permutation entropy averaged across participants. The same four functional brain atlases as in Extended Data Figure 4 were used (Brainnetome after Fan et al.^57^, Shen et al.^58^, Shirer et al.^59^, and the MMP after Glasser et al.^60^). The results from the main text are based on the Brainnetome atlas (gray background). The divergence between subcortical regions (high complexity) and cortex, especially in pericentral areas (low complexity), is equivalently observed with two volume-based atlases of higher (Shen) and lower (Shirer) spatial granularity as well as in surface-based extraction of cortical timeseries (Glasser).

**Extended Data Fig. 6.**
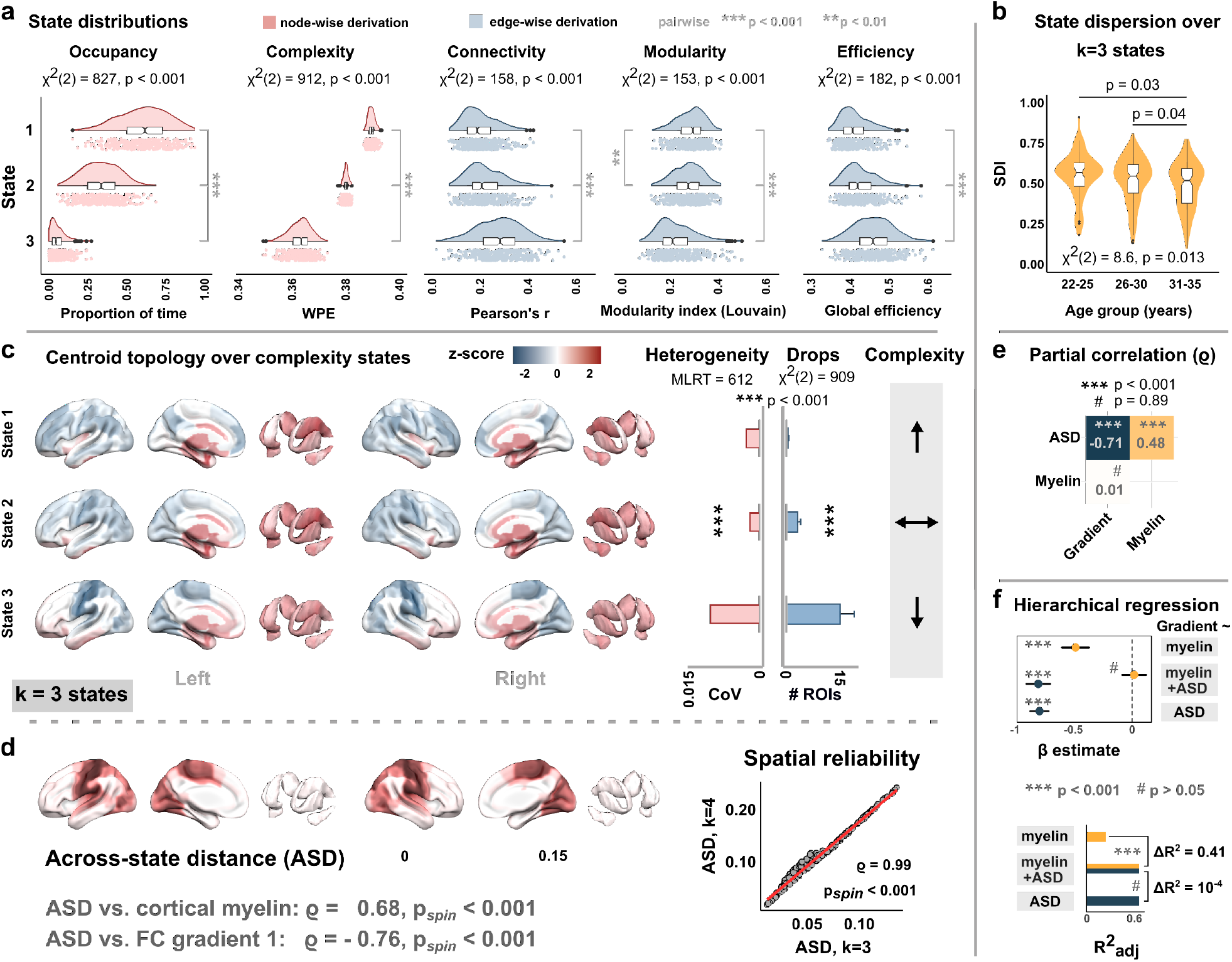
Validation analyses for k=3 expected complexity states. **a**, State-wise distributions of occupancy, signal complexity, FC strength, network modularity, and global efficiency. **b**, Age-related reduction of state exploration. **c**, Spatial topology, heterogeneity, and state-wise drop distribution over complexity states. **d**, Spatial topology, and spatial reliability of the across-state distance (ASD) compared to k=4 expected complexity states. **e**, Partial correlation between ASD, myelin, and the unimodal-to-transmodal connectivity gradient loadings. **f**, Hierarchical regression on the gradient loadings with myelin content and ASD as explanatory variables. All results closely follow the findings reported in Figures 3 and 4 of the main text.

**Extended Data Fig. 7.**
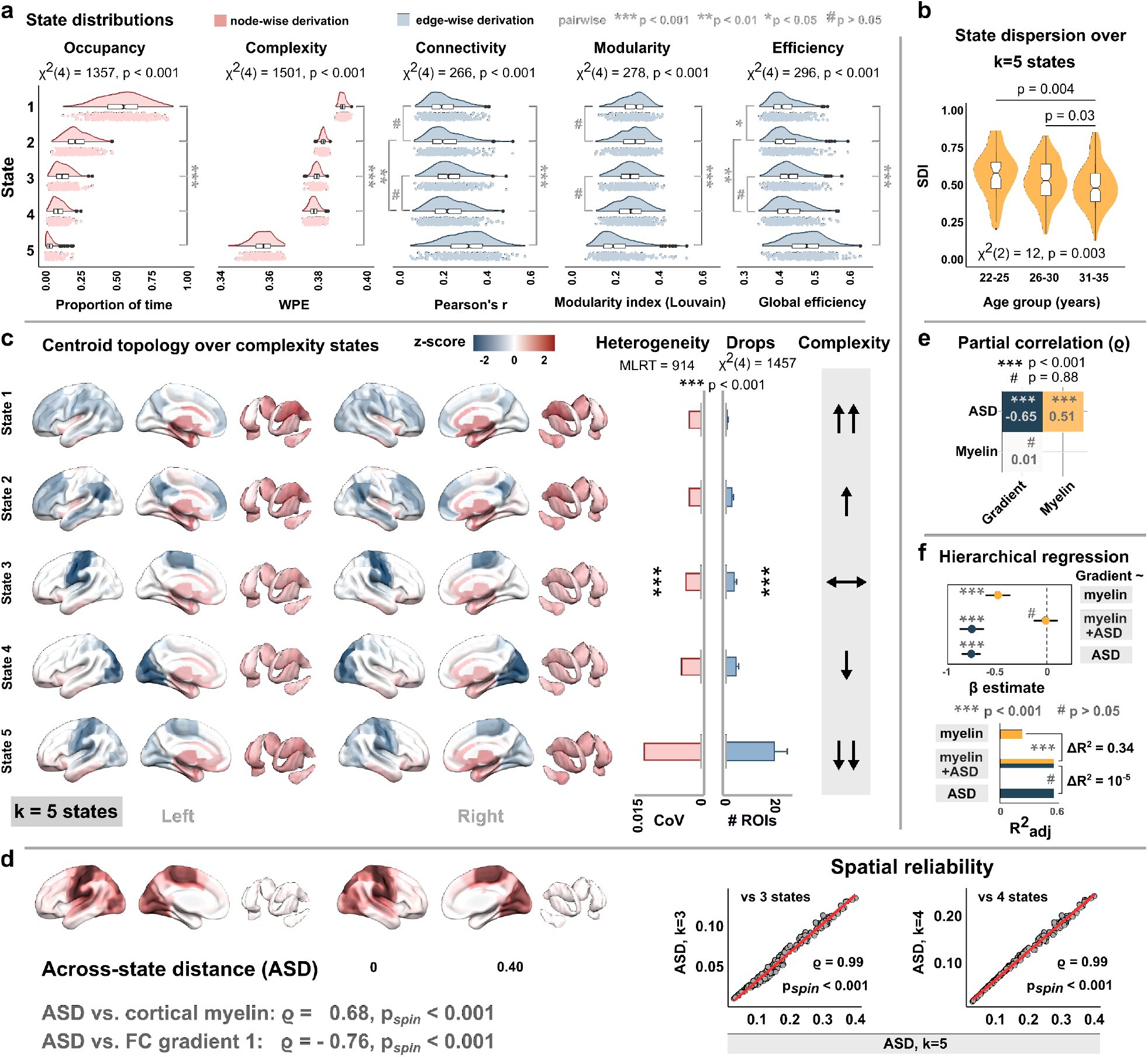
Validation analyses for k=5 expected complexity states. **a**, State-wise distributions of occupancy, signal complexity, FC strength, network modularity, and global efficiency. **b**, Age-related reduction of state exploration. **c**, Spatial topology, heterogeneity, and state-wise drop distribution over complexity states. **d**, Spatial topology, and spatial reliability of the across-state distance (ASD) compared to k=4 and k=3 expected complexity states. **e**, Partial correlation between ASD, myelin, and the unimodal-to-transmodal connectivity gradient loadings. **f**, Hierarchical regression on the gradient loadings with myelin content and ASD as explanatory variables. As for k=3 expected complexity states (Extended Data Figure 6), results closely follow the findings reported in Figures 3 and 4 of the main text.

**Extended Data Fig. 8.**
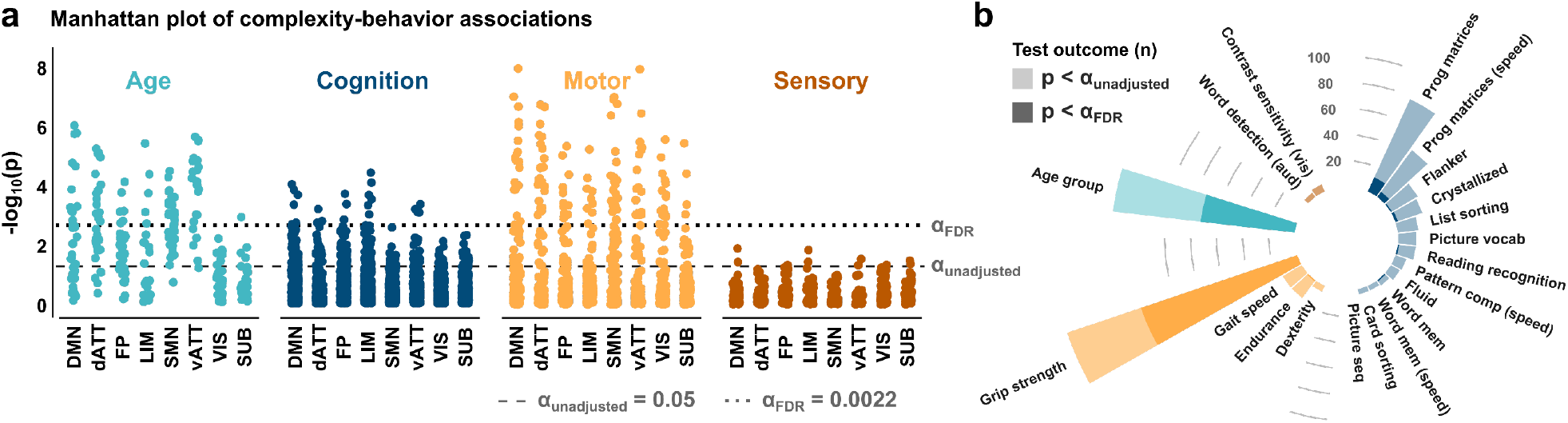
Univariate complexity-behavior associations over individual scores. **a**, Manhattan plot of negative log p-values for nonparametric correlation tests between behavioral variables and regional WPE values (dashed line: unadjusted alpha-level of 0.05; dotted line: critical alpha-level after FDR-correction over all tests). **b**, Test outcomes over the individual scores of age, cognition, motor function, and sensory task performance. To extract general behavioral features, dimensionality reduction through principal component analysis was applied to individual task performance metrics for the PLS analysis in the main text.

**Supplementary Movie 1. Visualizing complexity drops**. The clip shows the first 50 windows of regional complexity timeseries (cf., Fig. 1a), corresponding to ∼2.5 minutes of resting-state fMRI activity from an exemplary recording (HCP participant 11716, second scan). White indicates high-complexity activity of a given region, while colors indicate moderate (brown) to strong (blue) drops in complexity over the current window. Note the spread of complexity drops to increasingly more regions around window 10 (∼1 min) and 40 (∼2.1 mins), corresponding to individual drop cascades (Fig. 2a). These spreads are modelled with directed graphs (Fig. 2b) that underlie the reported propagation analyses (Fig. 2c-g).

